# ISG15 induces IL-10 production in human monocytes and is a biomarker of disease severity during active tuberculosis

**DOI:** 10.1101/166553

**Authors:** Paula Fernandes dos Santos, Johan Van Weyenbergh, Murilo Delgobo, Daniel de Oliveira Patricio, Brian J. Ferguson, Rodrigo Guabiraba, Tim Dierckx, Soraya Maria Menezes, André Báfica, Daniel Santos Mansur

**Affiliations:** Laboratory of Immunobiology, Department of Microbiology, Immunology and Parasitology, Universidade Federal de Santa Catarina, Campus Trindade, Centro de Ciências Biológicas, Bloco A, sala 213, Santa Catarina, Brasil. CEP 88040-900; Department of Pathology, University of Cambridge; ISP, INRA, Université François Rabelais de Tours, 37380 Nouzilly, France; Department of Microbiology and Immunology, Rega Institute for Medical Research, Laboratory for Clinical and Epidemiological Virology, KU Leuven - University of Leuven, Leuven, Belgium; Lead Contact

## Abstract

Interferon stimulated gene 15 (ISG15) deficiency in humans leads to severe interferonopathies and mycobacterial disease, the latter being previously attributed to its extracellular cytokine-like activity. Here, we demonstrate a novel role for secreted ISG15 as an IL-10 inducer, unique to primary human monocytes. Employing *ex vivo* systems analysis of human transcriptome datasets, we observed a significant correlation of ISG15-induced monocyte IL-10 and lymphocyte IFNγ expression. This effect was associated with p38 MAPK and PI3K signalling in healthy volunteers. The specificity and MAPK/PI3K-dependence of ISG15-induced monocyte IL-10 production was confirmed *in vitro* using CRISPR/Cas9 knockout and pharmacological inhibitors. Moreover, this *ISG15*/*IL10* axis was amplified in leprosy but disrupted in human active tuberculosis (TB) patients. Importantly, ISG15 strongly correlated with inflammation and disease severity during active TB. In conclusion, this study identifies a novel anti-inflammatory ISG15/IL-10 myeloid axis that is disrupted in active TB, revealing a potential biomarker for disease severity in this major human disease.

## Introduction

Type one interferons (IFN-I) exert most of their functions by inducing the expression of interferon-stimulated genes (ISGs). To date, over 300 ISGs have been described (de Veer et al., 2001; Der et al., 1998) and interferon stimulated gene of 15 KDa (ISG15) is prominently expressed in response to infection, in autoimmune diseases, cancer and physiological processes such as pregnancy (Dos Santos and Mansur, 2017; Hansen and Pru, 2014; Henkes et al., 2015; Hermann and Bogunovic, 2017; Tecalco Cruz and Mejia-Barreto, 2017; Wang et al., 2017a). ISG15 is synthesized as a 17 KDa precursor that is cleaved in the C-terminal region producing a mature form of 15 KDa. Also called ubiquitin cross-reactive protein (UCRP), ISG15 was the first ubiquitin-like protein to be described and it can be covalently linked to other proteins in a process called ISGylation (Dos Santos and Mansur, 2017; Haas et al., 1987; Loeb and Haas, 1992; Skaug and Chen, 2010). ISGylation is important for cell intrinsic immunity against several viruses including Influenza A, Vaccinia, Ebola, HIV and Hepatitis C virus (Morales and Lenschow, 2013; Schoggins and Rice, 2011; Skaug and Chen, 2010).

In addition to its intracellular ISGylation-mediated processes, the mature form of ISG15 can be secreted and possesses cytokine-like activities that modulate leukocyte functions (Bogunovic et al., 2013; Dos Santos and Mansur, 2017). For instance, soluble ISG15 was found to enhance production of IFNγ by lymphocytes and NK cells (Bogunovic et al., 2012; D'Cunha et al., 1996) and to stimulate NK cell proliferation (D'Cunha et al., 1996) as well as neutrophil migration (Owhashi et al., 2003). Importantly, ISG15 deficiency in humans is associated with a severe Mendelian susceptibility to mycobacterial disease (Bogunovic et al., 2012) and cells from patients with a nonsense mutation or a frame-shift in *isg15* are deficient in IFNγ-mediated immunity. This activity is attributed to the effects of extracellular ISG15 in NK cells and possibly occurs through an unknown receptor (Bogunovic et al., 2012). Furthermore, humans lacking ISG15 also develop exacerbated IFN-I-induced immunopathology (Zhang et al., 2015). This evidence suggests that extracellular or free ISG15, especially in humans (Speer et al., 2016), may regulate multiple aspects of the host immune response to pathogens and implicates this protein as an important component induced during infection and inflammatory processes involving IFN-I signalling. However, despite its ability to induce pro-inflammatory mediators, IFN-I may also exert anti-inflammatory effects (Billiau, 2006; Borden et al., 2007; McNab et al., 2015), and whether soluble extracellular ISG15 modulates anti-inflammatory responses has not been reported.

The present study demonstrates that ISG15 induces IL-10 synthesis in human primary monocytes through MAPK- and PI3K-dependent pathways. Additionally, analysis of human transcriptome data sets identified a myeloid *ISG15*/*IL10* axis present in homeostasis. In contrast, the *ISG15*/*IL10* axis is disrupted during active TB and *ISG15* mRNA levels strongly correlate with inflammatory and disease severity markers. These data suggest ISG15 may play a role in the crosstalk between Type I/Type II IFNs and IL-10 and reveal *ISG15* mRNA levels as potentially useful biomarker in human active TB.

## Results and Discussion

### ISG15 induces IL-10 production in human PBMC

Extracellular ISG15 stimulates IFNγ production by human NK cells (Bogunovic et al., 2012), so to investigate whether ISG15 regulates synthesis of other inflammatory cytokines, PBMCs were exposed to soluble ISG15 and 24h cell culture supernatants assayed for several cytokines by cytometric bead array (CBA). Out of this panel, only IL-10 and IL-6 were induced by ISG15 (Supplementary figure 1A). IL-10 is a key immune-regulatory cytokine that exerts opposing effects to IFNγ, hence this result was further assessed by treating PBMCs with different concentrations of pro- or mature ISG15 indicating a concentration-dependent response (Figure 1A). Following intracellular processing of pro-ISG15, its C-terminal LRLRGG domain is exposed and the protein becomes mature, a necessary requirement for ISGylation (Knight et al., 1988; Loeb and Haas, 1992; Narasimhan et al., 1996; Potter et al., 1999). Both pro- and mature ISG15 induced IL-10 secretion in human PBMCs in a similar manner (Figure 1A), indicating that LRLRGG sequence does not need to be exposed for ISG15-mediated IL-10 production. Exogenous ISG15 stimulated IL-10 synthesis by PBMCs from most of the healthy donors tested (Figure 1B). Control experiments showed that heat denatured ISG15 did not promote IL-10 synthesis demonstrating this protein requires its correctly folded structure to induce cell signalling (Figure 1B). Kinetic analysis of ISG15-stimulation in human PBMCs showed a peak of IL-10 mRNA and protein synthesis after 6 and 12 hours respectively (Figure 1C and D). Interestingly, this response was found to be specific for primary cells as a library of human cell lines (NKL, NK92, THP-1, Karpas, U937 and Jurkat) treated with ISG15 did not produce IL-10 (data not shown). Additionally, ISG15 treatment did not induce cell death by means of annexin V expression and propidium iodide (PI) incorporation (Figure 1E-G) suggesting IL-10 was actively secreted, not released from nor induced by apoptotic or necrotic cells. Together, these data show that ISG15 induces IL-10 synthesis and secretion by primary human PBMCs, independent of cell death.

**FIGURE 1.**
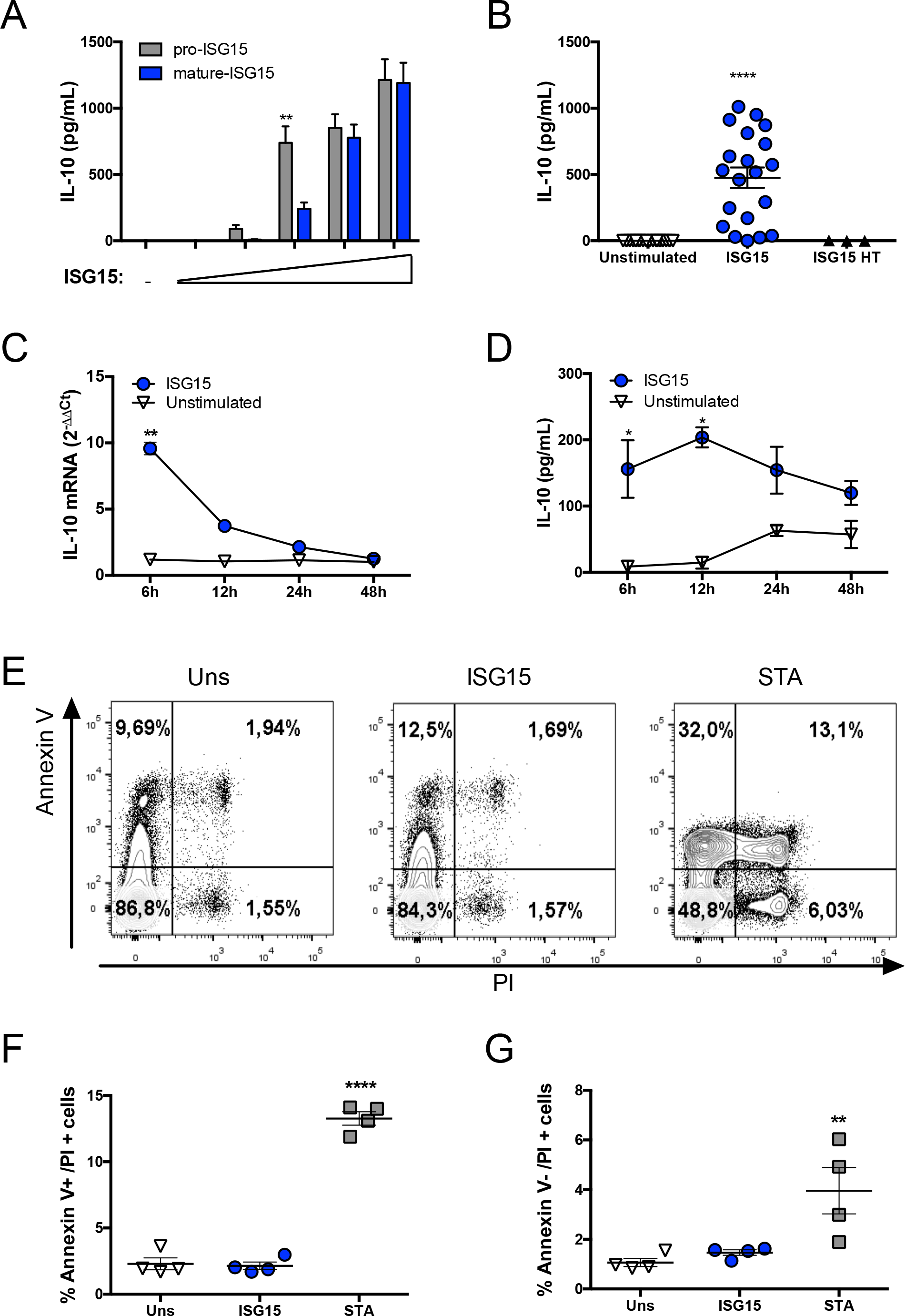
ISG15 induces the production of IL-10 in human PBMCs. (**A**) Dose-dependent IL-10 production as measured by ELISA 24 hours post stimulation of PBMCs with both pro- and mature ISG15 ([ISG15]: 0.15; 0.45; 1.5, 4.5 and 15 µg/mL). (**B**) Induction of IL-10 by recombinant, but not heat-treated, ISG15 using PBMCs from a total of 5 different donors in 8 independent experiments. (**C**) *IL10* mRNA expression in PBMCs treated with ISG15 at 6, 12, 24 and 48 hours post stimulation. (**D**) Quantification of IL-10 in the supernatant of human PBMCs at 6, 12, 24 and 48 hours after treatment with of ISG15 (**E**) Representative dot-plot evaluating cell-death in human PBMCs by annexin V and PI staining after treatment with ISG15 (2.0 µg/m) or Staurosporine (1 µM). (**F, G**) Quantification of cell death from the experiment described in (**E**). Unless stated otherwise, ISG15 concentration was 1.5 µg/mL. Error bars indicate SEM for biological replicates in each experiment. In each experiment PBMCs from 3 or more different donors were used. * P-value<0.05, ** p-value<0.01 and **** p-value<0.0001. ISG15HT, ISG15 heat-treated; STA, Staurosporine; PI, Propidium Iodide; Uns, Unstimulated.

### CD14^+^ cells are the main producers of ISG15-induced IL-10

ISG15 can act on different cell types (Bogunovic et al., 2012; D'Cunha et al., 1996; Owhashi et al., 2003; Recht et al., 1991) hence intracellular cytokine staining was used to identify the source of ISG15-induced IL-10 in PBMCs subpopulations. These experiments indicated that CD14^+^ cells are the main source of IL-10 (Figure 2A) with an average 2.5 fold-increase of CD14^+^IL-10^+^ cells as compared to unstimulated cultures (Figure 2B). Next, PBMCs were separated into CD14^+^ and CD14^−^ populations and both groups were exposed to soluble ISG15. Quantification of IL-10 and IFNγ 24 hours post stimulation confirmed the CD14^+^ population to be the main producers of IL-10 (Figure 2C) whilst we corroborated previous work showing the CD14^−^ population to be the source of ISG15-induced IFNγ (Figure 2D) (Bogunovic et al., 2012; D'Cunha et al., 1996). Additionally, these data indicate that recombinant ISG15-induced IL-10 synthesis by CD14^+^ populations does not require the presence of CD14^−^ cells.

**FIGURE 2.**
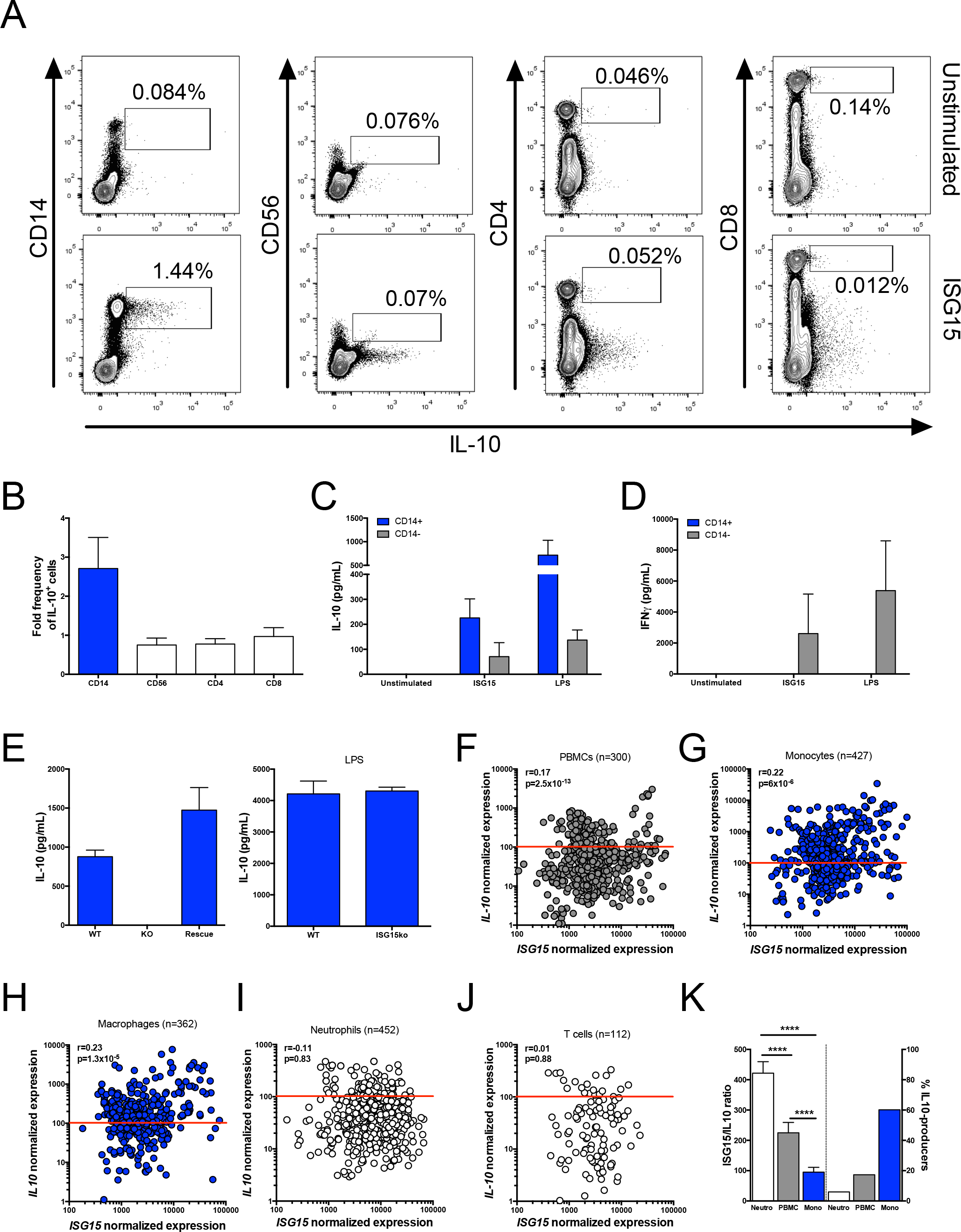
CD14^+^ cells are the main source of IL-10 upon ISG15 stimulation. (**A**) Representative dot plot of intracellular staining of IL-10 in CD14^+^, CD56^+^, CD4^+^ and CD8^+^ in ISG15-treated PBMCs. (**B**) PBMCs from 6 different individuals showing fold increase in IL-10 production from CD14^+^, CD56^+^, CD4^+^ and CD8^+^ populations after ISG15 stimulation. (**C-D**) ELISA quantification of IL-10 and IFNγ in the supernatants of CD14^+^ and CD14^−^ separated populations treated with ISG15 (**E**) A549 WT or ISG15 KO cells were co-cultured with primary CD14^+^ cells magnetically separated from PBMCs and IL-10 production was measured by ELISA 24 hours later. *ISG15* KO cells were also transfected with a plasmid expressing *ISG15* in order to rescue its function. LPS was used as a positive control for IL-10 stimulation. Error bars indicate SEM for biological replicates in each experiment. All experiments were repeated at least two times. In every experiment PBMCs from 3 or more donors were used. (**F-J**) Transcriptome datasets of healthy controls (ImmuCo, ImmuSort) confirm *ISG15* and *IL10 ex vivo* expression levels are strongly and positively correlated in total PBMCs (**F**), purified primary monocytes (G) and macrophages (**H**), but not neutrophils (**I**) or T cells (**J**). Red lines indicate the approximate threshold for *IL10* mRNA detection (determined for each individual microarray). (**K**) Neutrophils display the largest *ISG15*/*IL10* ratio *ex vivo*, whereas monocytes are the major *IL10* expressing leukocyte population (>PBMCs>neutrophils) under homeostatic conditions. **** p-value<0.0001.

To test whether endogenously produced ISG15 stimulates IL-10 synthesis, a co-culture experiment was set up using a lung epithelial cell line, A549, as a source of ISG15 (Wang et al., 2017b). For these assays, an *ISG15*-knockout (KO) A549 cell line was generated using CRISPR/Cas9 technology (Supplementary Figure 1B, clone 3). Wild type (WT) or *ISG15*-KO A549 cells were then co-cultured with purified human primary CD14^+^ cells or stimulated with LPS as a positive control. In this setup, A549-monocyte co-cultures led to a consistent production of IL-10, an outcome completely abrogated when *ISG15*-KO A549 cells were used. This effect could be rescued by re-introduction of the *ISG15* gene into the knockout cells (Figure 2E) thus demonstrating the specificity of epithelial cell-derived ISG15 for the induction IL-10. To study co-regulation of *ISG15*/*IL10*/*IFNG* pathways in different cell types *ex vivo*, we next examined transcriptome datasets of purified major human leukocyte subsets (ImmuCo, ImmuSort) (Wang et al., 2015a; Wang et al., 2015b). Expression levels of *ISG15* and *IL10* are positively correlated in total PBMCs, purified monocytes and macrophages, but not neutrophils and T-cells (Figures 2F-J). Although neutrophils display the highest *ISG15*/*IL10* expression ratio, monocytes are the main *ex vivo IL10* expressing cell type (60.19% of cells with IL10 transcripts above detection limit, vs. 17.33% in PBMCs and 5.97% in neutrophils, Fig. 2K), thus corroborating with our *in vitro* results (Fig. 2A-D). Consistent with a previous study (Tamassia et al., 2013), low or undetectable *IL10* transcripts in human neutrophils (Fig. 2K) are explained by the inactive chromatin configuration of the *IL10* locus in these cells. Together, this set of results suggests a role for extracellular rather than intracellular ISG15 as inducer of monocyte-derived IL-10 (this study) and NK-derived IFNγ (Bogunovic et al., 2012). Since the unique susceptibility of ISG15-deficient children to low virulence mycobacteria has underscored a role for extracellular ISG15 (Bogunovic et al., 2012), we performed a systems analysis approach to gain insights on the possible influence of ISG15/IL-10 axis during mycobacterial exposure in humans.

### IL-10 production in response to ISG15 requires MAPK-PI3K signalling pathways

Mitogen-activated protein kinase (MAPK) and phosphatidylinositol-4,5-bisphosphate 3-kinase (PI3K) signalling pathways have been shown to participate in *IL10* transcription in human monocytes and macrophages. For instance, p38, ERK1/2 and PI3K are crucial for IL-10 synthesis during microbial stimuli such as LPS and *Mycobacterium* (Ma et al., 2001; Nair et al., 2009). Thus, we analysed a published transcriptome dataset of latent TB using WebGEstalt and Ingenuity Pathway Analysis (IPA). As shown in Supplementary Table, MAPK and PI3K signalling pathways were significantly enriched in latent TB transcriptomes, as compared to healthy, uninfected controls. We next investigated whether members of MAPK and PI3K signalling families displayed divergent expression patterns between latent and active TB. MAPK family members were up-regulated in both latent and active TB (Figure 3A). However, *PIK3CA* (PI3Kalpha) levels were up-regulated in active TB and down-regulated in latent TB. In this scenario *MAPK14* (p38) expression levels were significantly and positively correlated to *ISG15* levels *ex vivo* (Figure 3B). Additionally, *MAPK3* (MAP3K/ERK1) levels were positively correlated to *IL10* (Figure 3C) and negatively correlated to *IFNG* transcript levels (Figure 3D). Finally, *PIK3CA* and *PIK3CB* (PI3Kbeta) transcripts were negatively correlated to *IL10* expression levels (Figure 3E). These data suggested that ISG15/IL-10 as well as MAPK/PI3K-associated transcripts are co-regulated during mycobacterial stimulation *in vivo* and raised the possibility that MAPK/PI3K pathway is involved in ISG15-induced IL-10 responses in monocytes. Indeed, following exposure of CD14^+^ cells to ISG15, increased phosphorylation of p38 MAPK was observed (Figure 3F). More importantly, the use of two distinct inhibitors for p38 (Figures 3G-H) as well as inhibitors for MEK1/2 (Figure 3I) and PI3K (Figure 3J) abrogated IL-10 production in ISG15-stimulated monocytes. In contrast, chloroquine, an inhibitor targeting DNA-PKCs/TLR9/endosome signalling pathways, did not affect IL-10 synthesis induced by ISG15 (Figure 3J). These results suggest a central role for p38 activation and MAPK as well as PI3K signalling in ISG15-induced IL-10 production by primary monocytes.

**FIGURE 3:**
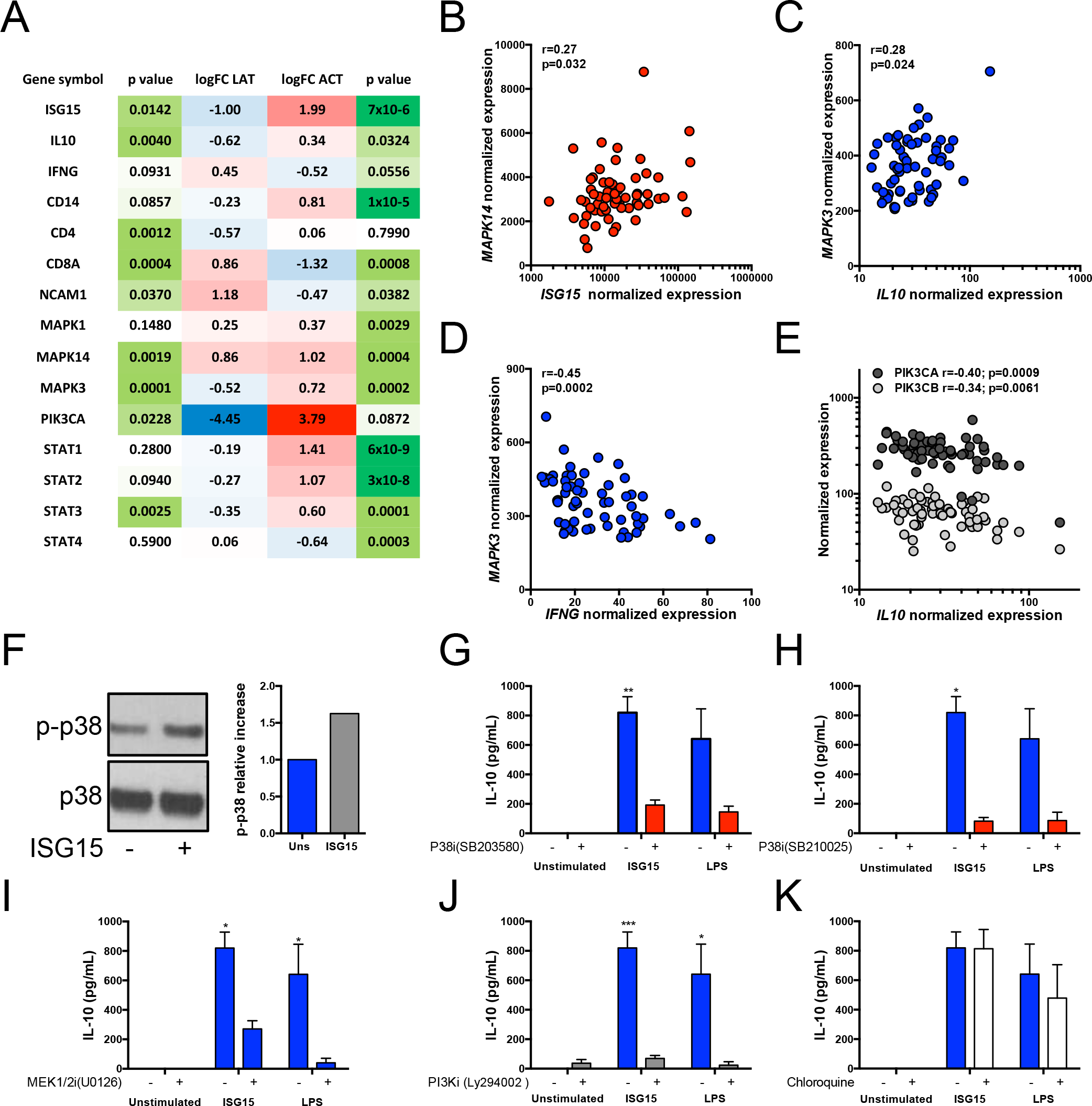
ISG15 induces monocyte derived IL-10 via p38, MEK1/2 and PI3K signalling pathways, which are deregulated in human mycobacterial infections. (**A**) MAPK family members expression in both latent and active TB. *Ex vivo* expression levels correlation of *MAPK14* and *ISG15* (**B**), *MAPK3* and *IL-10* (**C**), *MAPK3* and *IFNG* (**D**) and of *PIK3CA* or *PIK3CB* with *IL-10* (**E**) during latent TB infection (**F**) Representative immunoblot showing the phosphorylation of p38 MAPK 15 min after the stimulation of ISG15 in CD14^+^ cells. (**G-J**) CD14^+^ cells were treated for 1h with p38 (10 µM), MEK1/2 (50 µM) and PI3K inhibitors (50 µM) (B-E respectively) prior to addition of ISG15 (1µg/mL) or LPS (100 ng/mL). (**K**) Chloroquine (5µg/mL) was used as an unrelated control drug. 24 hours after treatment, supernatant was harvested and used for IL-10 quantification by ELISA. Error bars indicate SEM for biological replicates in two independent experiments. * P-value<0.05, ** p-value<0.01 and *** p-value<0.001. Uns, Unstimulated.

### An ISG15/IL10/IFNG cluster in healthy controls is disrupted during active TB

Altogether, our data indicated that ISG15 is associated with immunoregulatory responses and it could have an important role in mycobacterial-induced inflammation. To test this concept, publicly available transcriptome data sets from established cohorts of healthy controls and patients with leprosy as well as latent or active tuberculosis were examined. Positive and negative correlations (Spearman Rho) were calculated between normalized transcript levels of MAPK/PI3K/STAT signalling family members, established myeloid lineage markers (*CD14*, CD16=*FCGR3A/FCGR3B*, *ITGAM*, *ITGAX*) and lymphoid lineage markers (*CD4*, *CD8*, CD56=*NCAM1*, *ITGAL*) plus *ISG15*, *IL10* and *IFNG* transcripts. Unsupervised hierarchical clustering of these transcripts was then performed, based on the resulting correlation matrices. In healthy controls (Figure 4A), *ISG15* mRNA strongly clusters with *IL10*, and to a lesser extent with *IFNG*, indicating the existence of a regulatory balance between pro-and anti-inflammatory effects under homeostatic conditions. Leprosy lesions have been shown to express both type I IFN and IL-10, a scenario that leads to suppression of IFNγ effector activities (Teles et al., 2013). An unsupervised hierarchical cluster analysis of the cohort published by Teles and colleagues shows the myeloid anti-inflammatory *ISG15*/*IL10* axis maintained in leprosy lesions (Figure 4B), demonstrated by a single expression cluster comprised of, *ISG15*, *IL10* and monocyte (*CD14*) and myeloid markers (CD64=*FCGR1*, CD11c=ITGAX, PU.1=SPI1, CD16=FCGR3). Consistent with previous data (Teles et al., 2013), this disease cluster was negatively correlated to a “protective” *CD8*/*IFNG*/*STAT4* cluster, which is associated with milder (borderline tuberculoid/paucibacillary) clinical form, whereas the *ISG15*/*IL10*/*CD14* cluster was associated with the severe (lepromatous/multibacillary) disease form. Surprisingly, the *ISG15*/*IL10* axis was disrupted in whole blood transcriptomes of active TB (Figure 4C). However, ISG15 (but not *IL10* or *IFNG*) retained its association with disease status and monocyte/myeloid markers (*CD14, FCGR1*). Since TB disease signature in the whole blood is predominated by neutrophils (Berry et al., 2010) and monocytes only make up a minor fraction in these samples, we next investigated whether components of the *ISG15*/*IL10* signalling pathway might be overexpressed in purified monocytes from TB patients, as compared to control monocytes. Indeed, Ingenuity Pathway Analysis identified MAPK signalling as significantly enriched in monocytes from TB patients (Supplementary Table), and p38 MAPK was significantly interconnected with several TB signature genes (*FCGR1, IL27, SIGLEC6*) in a disease network (Figure 4D). Together, these results show the *ISG15/IL10* axis is disrupted during active TB.

**FIGURE 4:**
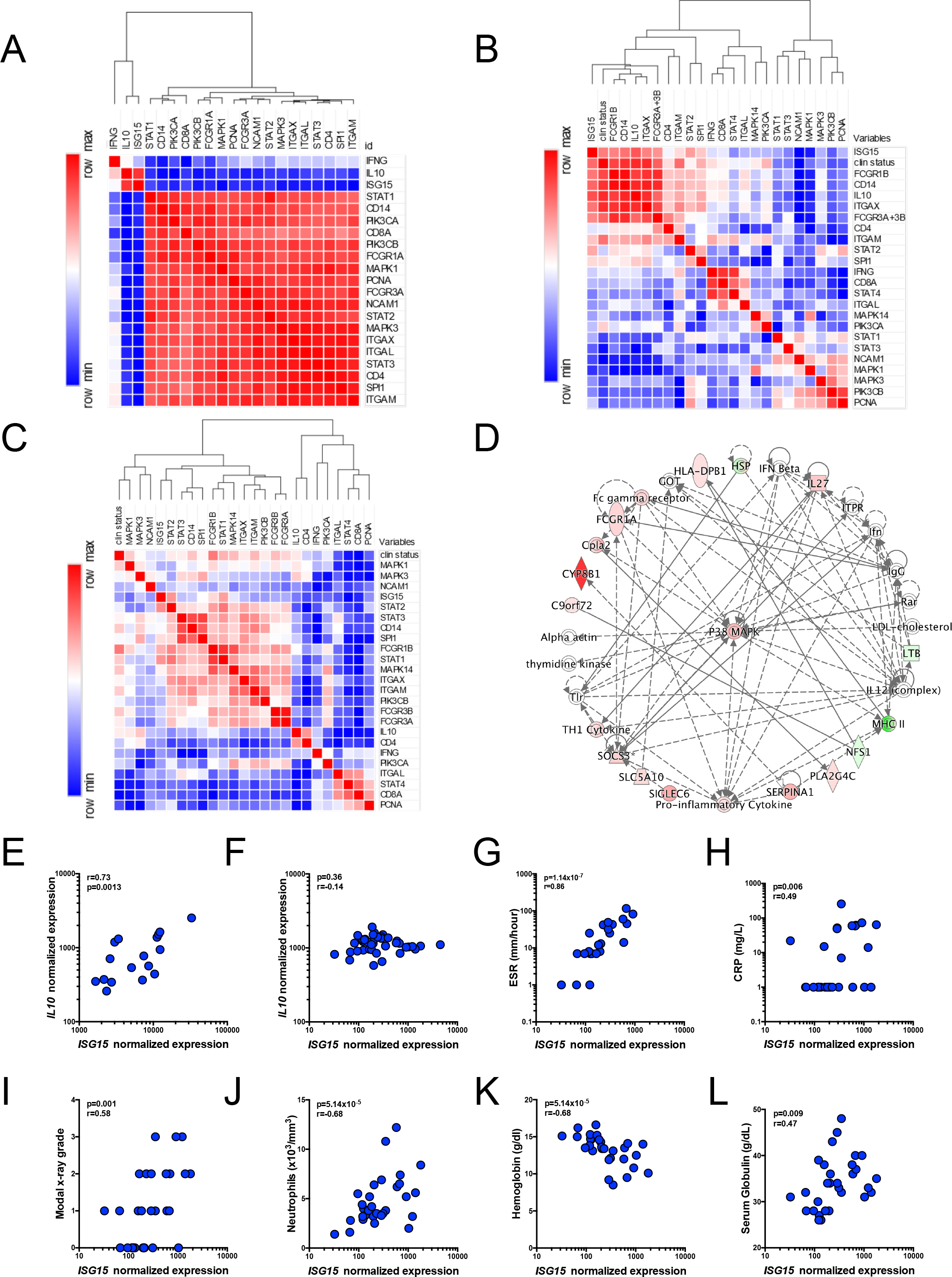
An anti-inflammatory *ISG15*/*IL10* myeloid axis is amplified in human leprosy and disrupted in human tuberculosis, revealing a novel clinical biomarker. (**A-C)** Heatmaps representing positive (red) and negative (blue) correlation matrix of selected genes (see text) classified by unsupervised hierarchical clustering (Eucledian distance). (**A**) Healthy controls (GSE80008) (**B**) Leprosy patients (GSE82160) (**C**) TB cohort (GSE85487) (**D**) Significantly enriched network (Ingenuity Pathway Analysis) showing p38 MAPK as highly interconnected in the monocyte transcriptome of TB patients. (**E-F**) *ISG15* transcript correlates with (**G, H**) established inflammatory metrics (erythrocyte sedimentation rate (ESR), C-reactive protein (CRP), (**I**) tissue damage (Modal X-ray grade) as well as (**J-L**) systemic clinical parameters (neutrophil count, haemoglobin, and globulin serum concentration).

To further investigate the possible connection between ISG15 and *M. tuberculosis*-mediated immunopathology, we examined the expression of this gene in an independent large cohort in which both detailed clinical parameters and corresponding transcriptome data are available (Berry et al., 2010). Expression levels of *ISG15* were significantly correlated with established inflammatory biomarkers such as erythrocyte sedimentation rate (ESR), C-reactive protein (CRP), tissue damage (Modal X-ray grade) as well as systemic clinical parameters (neutrophil count, haemoglobin, and globulin serum concentration) (Figure 4 G-L). These results suggest ISG15 may be a biomarker of disease severity in active TB patients.

Currently, there are no reports on the expression of ISG15 during human mycobacterial diseases *in vivo*. In the bioinformatics approach performed here, some but not all cohorts showed differences in ISG15 mRNA expression between healthy volunteers (HV) and acute TB patients (data not shown). However, since HV displayed high levels of ISG15 expression, it would be important to obtain parametric data from pre vs post-infection patients’ samples. While ISG15 is critical for IFNγ production by cells from vaccine strain *BCG*-infected patients (Bogunovic et al., 2012), this protein has a synergistic effect when combined with IL-12 (Bogunovic et al., 2012), an important inducer of IFNγ (Chan et al., 1992; Chan et al., 1991). Furthermore, IL-10 inhibits production of IL-12 and, consequently, of IFNγ by PBMCs (D’Andrea et al., 1993), pointing to a pleiotropic effect for ISG15. Cell type and context dependent effects of ISG15 could explain these diverse activities. This work and others (Bogunovic et al., 2012) suggest that despite ubiquitous expression in different cell types, neutrophils are a major source of secreted ISG15. Additionally, Mtb-infected macrophages can release microparticles containing ISG15 *in vitro* (Hare et al., 2015). Although we have not tested this directly, we speculate that soluble phagocyte-derived ISG15 is important to the orchestration of immune responses *in vivo*, driving the production of at least two major cytokines, IL-10 and IFNγ (Supplementary Figure 2). Interestingly, the intra and extracellular location of ISG15 and its ability to induce a plethora of effects in distinct cells, resembles the function of an alarmin (Rider et al., 2017). As shown in figure 4, ISG15’s function is context dependent, varying from a driver of an anti-inflammatory monocytic/IL-10 axis in homeostasis and the less severe *M. leprae* infection to a strong pro-inflammatory IFNγ-biased scenario during active TB. Additionally, since ISG15 and IL-10 lack correlation in active *M. tuberculosis* infection, it is possible that IL-10 is controlled by different signals other than ISG15 during disease. Whether virulent *M. tuberculosis* hijacks the ISG15/IL-10 axis contributing to induction of tissue pathology remains to be determined.

In conclusion, these findings confirm and extend previous work characterizing soluble extracellular ISG15 as a pleiotropic cytokine (or alarmin) that induces both pro- and anti-inflammatory effects in a variety of cell types. Moreover, the combined *ex vivo* and *in vitro* approach uncovers a novel myeloid *ISG15*/*IL10* p38-mediated anti-inflammatory signalling cascade, which is preserved in human leprosy but disrupted in active TB. Strikingly, our data indicate *ISG15* mRNA as a novel biomarker of disease severity during acute TB that merits further investigation.

## Supplemental figure 1

(**A**) ISG15 induces the production inflammatory cytokines. PBMCs from healthy donors were treated with ISG15 (15 µg/mL) and supernatant was harvested 24 hours for inflammatory cytokines quantification by cytometric Bead Array (CBA). ISG15 induced the production of IL-10, IL-6 and IL-1β. (**B**) Generation of ISG15 deficient A549 cell line. ISG15 deficient A549 cell line was produced using CRISPR/Cas9. After clone selection, cells were stimulated with IFNβ (1000 IU/mL), proteins extracted after 24 hours and immunoblotted with anti-ISG15 and anti-tubulin antibodies. Clone 3 (ISG15 KO) was used for further experiments.

## Supplemental figure 2

Proposed model. ISG15 induces both IL-10 (Blue) and IFNγ (Red) biased responses in humans. The ISG15/IL-10 myeloid axis is present healthy individuals and also in leprosy lesions while ISG15/IFNγ lymphoid axis is characteristic of anti-Mtb immunity but is also related to immunopathology, a state in which ISG15 transcripts strongly correlates with disease severity parameters when the myeloid axis is disrupted. Dotted lines represent known literature (Chomarat et al., 1993; Redford et al., 2011).

## Supplemental Table

Tab 1. PI3K gene expression enriched during latent tuberculosis.

Tab 2. MAPK gene expression enriched during latent tuberculosis.

Tab 3. Ingenuity Pathway Analysis for MAPK in monocytes during latent tuberculosis.

Tab 4. p38 network in monocytes during active tuberculosis.

## Material and methods

### Reagents

ISG15 was purchased from Boston Biochem and tested for endotoxins by R&D Systems (endotoxin value for lot #DBHF0614021 is <0.00394 EU/µg). LAL assay (Lonza) was performed according to the manufacturer's instruction and the endotoxin level of recombinant ISG15 was below the detection threshold. Pro-ISG15 (UL-615) was also purchased from Boston Biochem. *E. coli* LPS (strain O111:B4) (Invivogen) was used as a positive control for IL-10 production in human PBMCs and monocytes. P38 kinase inhibitors SB203580 and SB220025 (Calbiochem) were used at 10 µM, MEK1/2 inhibitor U0126 (Cell Signaling) at 50 µM and PI3K inhibitor Ly294002 (Cell Signaling) was at 50 µM. Solvent (DMSO, medium) was used as negative control and chloroquine (Sigma, 5µg/mL), a DNA-PKC/TLR endosomal signalling inhibitor, was used as an additional negative control.

### Primary human cells

Human PBMCs were separated from healthy individuals using Ficoll-paque (GE) according to manufacturer's instructions. Briefly, blood was collected in heparin-containing tubes, and gently mixed 1:1 with saline solution and gently mixed before being added over one volume of Ficoll-paque reagent. The gradient was centrifuged for 35 minutes at 400 × g, 18°C. PBMCs were harvested and washed once with 45 mL of saline solution for 10 min at 400 × g, 18°C. Subsequently, cell pellet was suspended and washed twice with 5 mL of saline solution for 10 min at 200 × g, 18°C to remove platelets. The remaining cell pellet was suspended to the desired density in RPMI 1640 (Gibco) supplemented with 5% foetal calf serum (Hyclone), 2mM L-glutamine (Gibco), 1 mM sodium pyruvate (Gibco), 25 mM HEPES (Gibco), 100 U/mL penicillin and 100 µg/mL streptomycin (Gibco). Cells were plated as described in each experiment. Human primary monocytes (CD14^+^ cells) were separated from PBMCs using CD14 microbeads (Miltenyi Biotec) according to manufacturer's instructions with the exception of the MACS buffer, which was prepared using 3% foetal calf serum. Monocyte enrichment varied between 73 to 92% between experiments. The use of PBMCs from healthy donors was previously approved by UFSC ethical committee (IRB#283/08).

### Generation of isg15 knockout cell lines

A549 lung epithelial cells were co-transfected with three gRNA/Cas9/GFP plasmids (provided by Horizon) targeting the *ISG15* locus using JetPEI (PolyPlus Transfection). The guide RNAs used were 5′ GGCTGTGGGCTGTGGGCTGT 3′, 5′ GGTAAGGCAGATGTCACAGG 3′ and 5’ TGGAGCTGCTCAGGGACACC 3’.72 hours after transfection, cells sorted for GFP fluorescence and then separated by limiting dilution. Single-cell derived clones were selected for ISG15 expression (Supplementary Figure 1B).

### A549 – CD14+ co-culture

A549 WT or ISG15 KO cells were seeded at 2 × 10^5^ cells/ml in 24 well-plates. Cells rested in the incubator for 6 hours before ISG15 KO cells were transfected with ISG15-pCEP4 plasmid using FugeneHD reagent (Promega) according to manufacturer's instructions. Cells were then washed and LPS was added 18 hours after transfection and immediately prior to the addition of a 2 × 10^5^ CD14^+^ cells overlay. Following 24 hours of co-culture, supernatants were harvested for IL-10 quantification.

### Immunoblotting

1 × 10^5^ CD14^+^ cells were added to a 96-well plate and left to rest overnight. Cells were stimulated with ISG15 (1µg/ml) and after 15 minutes cells were spun at 4°C, supernatant was removed and M-PER lysis buffer (Thermo Scientific) containing protease inhibitors (Complete, Mini Protease Inhibitor Tablets, Roche) and phosphatase inhibitors (#524625, Calbiochem) was added to the cells. Protein separation was performed according to M-PER manufacturer's instructions. Antibodies concentrations for detection of p38 (Cell Signaling #9212) and p-p38 (Cell Signaling #9211), ISG15 (Cat: A600, R&D Systems) and anti-α-Tubulin (clone DM1A, Millipore) were those suggested by the manufacturers. For Western blots, at least 20 µg of total protein were separated and transferred to a PVDF 0.22 µm blotting membrane. Membrane was blocked for at least 1 hour with 1X Tris Buffered Saline-0.1% Tween20 (TBST) with 5% w/v non-fat dry milk and subsequently washed 3 times with TBST for 5 minutes each wash. Membrane was incubated with primary antibodies diluted in 5% w/v BSA, 1X TBS, 0.1% Tween20 at 4°C with gentle shaking overnight. Membrane was washed 3 times for 5 min each with TBST and then incubated with the appropriate secondary HRP-linked antibody for 1 hour at room temperature. Membrane was washed 3 times of 5 min each with TBST before detection with ECL chemiluminescent substrate (Pierce).

### p38 MAPK and PI3K signalling pathway inhibition

1 × 10^5^ CD14^+^ cells were added to a 96-well plate and left to rest overnight. Inhibitors were added to cells for 1 hour prior to ISG15 treatment. 24 hours after treatment, cells were spun at 4°C; supernatant was harvested and IL-10 was quantified by ELISA.

### Cytokine quantification

For exploratory experiments, IL-1β, IL-6, IL-10, IL-12p70 and TNF were quantified in supernatants by human inflammatory cytokine cytometric beads array kit (CBA, BD Biosciences). IL-10 and IFNγ were quantified using Human IL-10 DuoSet ELISA kit (R&D Systems) or Human IFNγ mini kit (Thermo Scientific) according to manufacturer's instructions.

### Flow Cytometry Assays

PBMCs were seeded at a density of 5 × 10^5^ cells per well in 150 µL of medium. After 8 hours of resting at 37°C with 5% CO_2_, cells were treated with ISG15 (2 µg/mL) or LPS (1 µg/mL), unless indicated otherwise. Golgi Plug protein transport inhibitor (BD Biosciences) was added 1-hour post treatment, according to manufacturer's instructions. Then, 12 hours post treatment; growth medium was removed and cold 1X HBSS (Gibco) with 2.5 mM EDTA was added to each well. The tissue culture dish was kept at 4°C for 30 minutes and cells were suspended and transferred to 1.5 mL tubes. Cells were washed in a final volume of 1mL cold 1X HBSS (Gibco) with 2.5 mM EDTA at 300 × g and 4°C for 5 minutes. Supernatant was removed and cells were suspended in FACS buffer (1% BSA, 1% sodium azide in 1X PBS). Anti-human antibody mix containing anti-CD4 APC-Cy7 (clone OKT1) (BioLegend), anti-CD8 PE-Cy7 (clone SK1) (BioLegend), anti-CD14 PerCP-Cy5.5 (clone M5E2) (BioLegend), anti-CD56 FITC (clone NCAM 16.2) (BD Biosciences) was added to the cell suspension for 40 min at 4ºC in the presence of 10% AB blood-type human serum to block Fc receptors. Afterwards, cells were washed once with 1 mL of 1X PBS at 300 × g, 4°C, for 5 minutes and 1 mL of fixation buffer (1% paraformaldehyde in 1X PBS) was added to the cells. Tubes were kept in the dark at room temperature for 15 min and then centrifuged at 300 × g, 4°C, for 10 min to remove supernatant. 1 mL of permeabilization buffer (0.5% saponin in FACS buffer) was added to the cells and tubes were centrifuged at 300 × g, 4 C for 10 min. Intracellular stain with anti-IL-10-PE (clone JES3-9D7) (BioLegend) was carried out for 30 min in the dark at room temperature. Cells were washed with permeabilization buffer at 300 × g, 4°C, 5 min, supernatant was removed and cells were suspended in FACS buffer prior to acquisition of 1 × 10^5^ events or more. For analysis, all acquired events displayed as forward scatter (FSC) and side scatter (SSC) parameters were selected. After that, single cells events were selected using FSC area and height parameters (FSC-A × FCS-H) and auto-fluorescence was excluded using APC as an open channel. Intracellular IL-10 was then quantified in monocytes (CD14^+^IL-10^+^), NK cells (CD56^+^IL-10^+^), CD4 (CD4^high^IL-10^+^) and CD8 T cells (CD8^high^IL-10^+^). Gates were set according to unstained PBMC sample and controls. All samples were acquired on a Becton-Dickinson Verse flow cytometer using BD FACSuite^TM^ software. In order to analyse cell death, 5×10^5^ PBMC/well were treated with ISG15 2 µg/mL or Staurosporine (Sigma) 1 µM for 24 hours. Cells were then harvest, washed with 1 mL of PBS at 300 × g, room temperature, 5 min, supernatant was removed and cells were washed once in 1 mL of 1x Annexin biding buffer (eBioscience). Cell were resuspended at 10^6^ cells/mL in 1x Annexin biding buffer and FITC conjugated Annexin V (eBioscience) was added to the cell suspension for 15 min, room temperature, according to manufacture's instruction. Following incubation period, cells were washed with 1 mL of 1x Annexin biding buffer, 300 × g, room temperature, 5 min and resuspended in 200 uL of 1x annexin biding buffer. Propidium iodide (BD Pharmingen) was added at 0.25 µg/mL to the cell suspension prior to sample acquisition. Samples were acquired on a Becton-Dickinson Canto II flow cytometer using BD FACSDiva^TM^ software.

### Real time quantitative PCR (qPCR)

For relative quantification of *IL10* gene expression, total RNA was extracted from PBMCs treated or not with ISG15. RNA was extracted after 6, 12, 24 or 48 hours of treatment using RNeasy RNA extraction kit (Qiagen). Using 400 ng of RNA, cDNA was produced with High-Capacity cDNA Reverse Transcription Kit (Applied Biosystems) and 2µL of the product was used for the qPCR reaction in a final volume of 10 µL. qPCR reactions were performed using the primers forward 5'GAG ATC TCC GAG ATG CCT TCA G 3'and reverse 5'CAA GGA CTC CTT TAA CAA CAA GTT GT 3’ (Skrzeczynska-Moncznik et al., 2008). Fold-increase in *IL10* gene expression was determined by relative quantification using hypoxanthine phosphoribosyltransferase (*HPRT*) as endogenous control. Primers forward and reverse for *HPRT* were 5’ CCTGCTGGATTACATCAAAGCACTG 3’ and 5’ TCCAACACTTCGTGGGGTCCT 3’, respectively, and were used at 250 nM each.

### Microarray analysis

Curated and annotated publicly available data-sets (Berry et al., 2010; Novais et al., 2015; Speake et al., 2015; Teles et al., 2013; Wang et al., 2015a; Wang et al., 2015b)(GXB, ImmuCo, ImmuSort, BioGPS, GEO) were obtained from large, established cohorts of healthy controls, latent and active tuberculosis patients, comprising *ex vivo* and *in vitro* whole blood, total PBMCs, purified leukocyte subsets, non-leukocyte human primary cells and skin biopsies (leprosy patients, healthy controls and cutaneous leishmaniasis as non-mycobacterial infectious control). Novel datasets were generated for both whole blood and PBMCs from healthy controls and individuals infected with other non-mycobacterial intracellular pathogens (*Leishmania*, HIV-1, HTLV-1). PBMCs were isolated as above and immediately frozen in Trizol to preserve RNA integrity. Following Trizol extraction, total RNA was further purified using an RNeasy kit according to the manufacturer's protocol (QIAGEN, Venlo, Netherlands). Affymetrix Whole Genome microarray analysis was performed by the VIB Nucleomics Facility (Leuven, Belgium) using a GeneChip^®^ Human Gene 1.0 ST Array with the WT PLUS reagent kit (Affymetrix, Santa Clara, CA, USA) according to the manufacturer's specifications. Data preprocessing (RMA) was performed using the Bioconductor xps package. All microarray raw data are available at Gene Expression Omnibus database (GEO, http://www.ncbi.nlm.nih.gov/geo/) under series accession numbers GSE80008, GSE82160, GSE85487.

### Enrichment analysis

The Ingenuity Pathway Analysis (IPA) program was used to perform the initial pathway/function level analysis on genes determined to be differentially expressed in the microarray analysis (Ingenuity Systems, Red Wood City, CA). Uncorrected p-values and absolute fold-changes were used with cut-offs of p<0.05. Based on a scientific literature database, the genes were sorted into gene networks and canonical pathways, and significantly overrepresented pathways were identified. Further enrichment analysis was performed, including Gene Ontology (GO) term enrichment using the WEB-based GEne SeT AnaLysis Toolkit (WebGestalt), KEGG pathway enrichment using the pathway database from the Kyoto Encyclopedia of Genes and Genomes and transcription factor target enrichment using data from the Broad Institute Molecular Signatures Database (MSigDB). Genesets from the GO, KEGG pathways, WikiPathways and Pathway Commons databases, as well as transcription factors, were considered overrepresented if their corrected p-value was smaller than 0.05. Principal component analysis, correlation matrices (Spearman), unsupervised hierarchical (Eucledian distance) clustering were performed using XLSTAT and visualized using MORPHEUS (https://software.broadinstitute.org/morpheus/).

### Data processing and statistical analyses

Data derived from *in vitro* experiments was processed using Graphpad Prism 6 and analysed using unpaired Student's T test unless stated otherwise. Statistical significance is expressed as follows: * P-value<0.05, ** p-value<0.01, *** p-value<0.001 and **** p-value<0.0001. In all cases, data shown are representative from at least two independent experiments. Data from experiments performed in triplicate are expressed as mean ± SEM.

## Acknowledgements

DSM received support from CAPES Computational Biology (23038.010048/2013-27), CNPq Universal (473897/2013-0) and the Academy of Medical Sciences/UK (NAF004/1005). PFS and MD received CAPES and CNPq student fellowships respectively. AB received financial support from NIH-GRIP (TW008276), HHMI-ECS (55007412). AB is CNPq-PQ scholar and CAPES/ESE. BJF received support from an Isaac Newton Trust/Wellcome Trust ISSF/University of Cambridge research grant and a Wellcome Trust Seed Award (201946/Z/16/Z). TD received grant support from VLAIO (IWT141614). JVW received grant support from CAPES (PVE) and FWO (G0D6817N).

We would like to thank Prof. Aristobolo Mendes Silva from UFMG for providing reagents, suggestions and Dr. Alan Sher for critical reading and evaluation of these results.

The authors declare no conflict of interest.

## Author contributions

PFS, JVW and RG designed, performed experiments, analysed the data and wrote the manuscript; MD, DOP, TD and SMM performed experiments and analysed the data, BF, AB and DSM designed experiments analysed the data and wrote the manuscript.

